# Low cost, medium throughput depletion-binding assay for screening S-domain-receptor ligand interactions using *in planta* protein expression

**DOI:** 10.1101/2021.06.16.448648

**Authors:** Lin-Jie Shu, Milena Schäffer, Sabine Eschrig, Stefanie Ranf

## Abstract

**Background:** Plant cell-surface receptors sense various ligands to regulate physiological processes. Matching ligand-receptor pairs requires evidence of their direct interaction and is often a bottleneck in functional receptor studies. The S-domain-type (SD) pattern recognition receptor LORE senses medium chain-3-hydroxy fatty acids (mc-3-OH-FAs) such as 3-hydroxy decanoic acid (3-OH-C10:0) via its extracellular domain (ECD) They are perceived as signals of danger from Gram-negative bacteria and activate immune responses in *Arabidopsis thaliana*. LORE is found in low levels *in planta* and is poorly expressed in heterologous systems. Furthermore, chemical modifications of the mc-3-OH-FA ligand affect its biological activity. Taken together, this makes LORE-mc-3-OH-FA binding studies particularly challenging.

**Results:** To investigate the LORE-mc-3-OH-FA interaction, we have developed a sensitive assay system based on protein expression *in planta*. The ECDs of LORE and other proteins of interest were transiently expressed as soluble, apoplastic mCherry fusion proteins in *Nicotiana benthamiana* and collected in apoplastic washing fluids. Protein-ligand complexes and unbound ligand were separated according to their molecular weight. In a two-step procedure, we first investigated whether the ECD-mCherry fusion protein depletes 3-OH-C10:0 from the low molecular weight fraction (step ‘depletion’). Subsequently, protein-bound 3-OH-C10:0 retained in the high molecular weight fraction in the depletion step is released and detected (step ‘binding’). Both the unbound and the released 3-OH-C10:0 ligand are detected by a sensitive bioassay using *LORE* loss- and gain-of-function Arabidopsis plants. Using the depletion-binding assay, we show that the ECD of AtLORE and its ortholog from *Capsella rubella*, CrubLORE, bind 3-OH-C10:0. The ECD of AtSD1-23, the closest paralog of AtLORE, and mCherry did not bind 3-OH-C10:0 and are suitable negative controls.

**Conclusion:** The depletion-binding assay is a simple method for reliably detecting interactions between plant-expressed SD-type receptor ectodomains and mc-3-OH-FAs. It does not require special equipment or expensive consumables and is suitable for medium throughput screening. The assay is very flexible and can be easily adapted to investigate ligand interactions of other extracellular receptor domains.

## BACKGROUND

Plants possess an expanded and diversified repertoire of cell surface-localized receptor-like kinases (RLKs) and receptor-like proteins (RLPs) (Shiu et al., 2004) that function as receptors, co-receptors, scaffolds or regulators of receptor complexes involved in growth, development, response to abiotic stimuli, and immunity (Couto and Zipfel, 2016; Boutrot and Zipfel, 2017; Gou and Li, 2020). RLKs/RLPs bind chemically and structurally diverse ligands ranging from complex peptides, carbohydrates and lipids to low molecular weight hormones, nucleotides and metabolites via various extracellular domains (ECDs). According to their ECD composition, RLKs/RLPs are classified e.g. as leucine-rich repeat (LRR), Lysin-motif (LysM), S-domain (SD), Malectin-like domain (MLD) or L-type lectins (Lec) (Shiu and Bleecker, 2001; Boutrot and Zipfel, 2017; Dievart et al., 2020). To date, only a few RLKs/RLPs have been characterized in detail, among them several with a role in plant immunity. Pattern-recognition receptors (PRRs) sense molecular signatures characteristic of microbes or indicative of cellular damage, termed microbe- or damage-associated molecular patterns (MAMPs or DAMPs, respectively), and activate local and systemic immune responses referred to as pattern-triggered immunity (PTI) (Boutrot and Zipfel, 2017). The PRRs AtFLS2 and AtEFR, which sense the bacterial flagellin peptide epitope flg22 and elongation factor Tu epitope elf18, respectively, together with their common LRR-type co-receptors of the AtSERK family, are important prototype models of the LRR-RLK class. The LysM-RLK module AtCERK1-AtLYK5 from Arabidopsis and the LysM-RLP OsCEBiP together with the LysM-RLK OsCERK1 from rice, that mediate sensing of chitin oligomers from fungal cell walls, have also been studied in detail (Couto and Zipfel, 2016). Many more LRR- and LysM-RLKs/RLPs have been functionally characterized and 3D structures with bound ligands have been resolved for several LRR- and LysM-ECDs, making the LRR- and LysM-RLKs/RLPs the best-studied receptor classes to date (Couto and Zipfel, 2016; Hohmann et al., 2017). Overall, these studies have led to a fundamental understanding of cell-surface receptor signalling in plants. Although the number of RLKs/RLPs with a known function is steadily increasing, the identification of ligands and ligand binding studies often remain challenging.

Genetic studies, e.g. using RLK/RLP loss- or gain-of-function approaches, are instrumental in identifying potential ligand-receptor pairs, but additional methods are required to prove a direct interaction between a ligand and its receptor. Various different methods have been used to assess ligand binding to plant receptors (Sandoval and Santiago, 2020). The flg22-FLS2 interaction has been demonstrated, for example, by radioligand binding assay and chemical cross-linking using immunoprecipitated receptors from Arabidopsis and AtFLS2-exressing tomato cells (Chinchilla et al., 2006). Crystallization of the ECDs of AtFLS2 and its coreceptor AtSERK3 together with flg22 revealed the precise mode of the ligand-receptor interaction (Sun et al., 2013b). Chitin was also found to bind directly to AtCERK1 *in vitro* by radioligand binding assays (Iizasa et al., 2010) and by isothermal titration calorimetry (ITC) (Liu et al., 2012) using recombinant protein expressed in yeast and insect cells, respectively. According to ITC analyses, the LysM-RLK AtLYK5 binds chitin oligomers with higher affinity than AtCERK1 and forms a chitin-induced complex with AtCERK1 (Cao et al., 2014). AtDORN1, an L-type lectin receptor kinase that mediates the perception of extracellular ATP in Arabidopsis, binds radiolabeled ATP after recombinant expression in *Escherichia coli* bacteria (Choi et al., 2014). Many other techniques to study biomolecular interactions have been applied in plant receptor research, such as surface plasmon resonance (SPR), microscale thermophoresis (MST), and grating-coupled interferometry (GCI), all of which have their specific advantages and disadvantages (Sandoval and Santiago, 2020). While such methods allow quantification of binding properties, many of them require large amounts of purified receptor protein or ligand labelling, as well as specialized facilities, costly equipment or expensive consumables.

The class of the SD-RLKs has about 40 members in Arabidopsis (Shiu and Bleecker, 2003; Teixeira et al., 2018). Their ectodomains comprise a lectin domain, an epidermal growth factor-like (EGF) domain, and a plasminogen/apple/nematode (PAN) domain. Because of the similarity of their lectin domain to the *Galanthus nivalis* agglutinin (GNA), SD-RLKs are also called bulb-type or G-type lectin RLKs (Teixeira et al., 2018). The best-studied SD-RLKs to date are the S-locus receptor kinases (SRKs) that mediate self-incompatibility in Brassica. Haplotype-specific recognition of S-locus cysteine-rich (SCR) peptide (also called S-locus Protein 11, SP11) expressed on pollen by the stigma-expressed SRK leads to rejection of self-pollen to avoid inbreeding (Ivanov et al., 2010). The crystal structures of the ECDs of the *Brassica rapa* SRKs, SRK9 and SRK8, in complex with their ligands, SCR9 and SCR8, respectively, have been resolved and shown to form 2:2 SRK:SCR heterotetramers (Ma et al., 2016; Murase et al., 2020). Other SD-RLKs from, e.g. Arabidopsis, rice, tomato, wild soybean, and wheat are involved in plant-microbe interactions, abiotic stress responses, development and growth, but their ligands remain mostly unknown (Charpenteau et al., 2004; Chen et al., 2006; Bonaventure, 2011; Chen et al., 2013; Cheng et al., 2013; Sun et al., 2013a; Zhao et al., 2013; Catanzariti et al., 2015; Ranf et al., 2015; Zou et al., 2015; Fan et al., 2018; Pan et al., 2020). The *A. thaliana* SD-RLK LIPOOLIGOSACCHARIDE-SPECIFIC REDUCED ELICITATION (AtLORE), alias SD1-29, was initially identified in a forward genetics screen for mutants that are insensitive to treatment with lipopolysaccharide (LPS) extracts from Pseudomonas bacteria (Ranf et al., 2015). Recently, we showed that medium chain-3-hydroxy fatty acids (mc-3-OH-FAs) with eight to twelve carbons activate LORE-dependent immune responses. Free 3-OH-C10:0 triggered the strongest response among the immunogenic mc-3-OH-FAs and chemical modifications decreased its eliciting activity with increasing complexity (Kutschera et al., 2019). Free 3-OH-C10:0 was found in the bacterial LPS extracts. Highly purified LPS, that did not contain detectable amounts of 3-OH-C10:0, did not activate immune responses in Arabidopsis (Kutschera et al., 2019). Thus, LORE is a PRR that senses mc-3-OH-FAs.

We have previously shown that 3-OH-C10:0 binds *in vitro* to the ECD of AtLORE (eAtLORE) recombinantly expressed as secreted protein in insect cells (Kutschera et al., 2019). However, the yield of eAtLORE protein from insect cells was low and relatively high salt concentrations were required to keep the protein in solution. The stability and function of plant proteins recombinantly expressed in heterologous systems such as insect cells, yeast or bacteria may be affected by misfolding or, in the case of extracellular proteins, by differential or absent glycosylation. Heterologous expression of the ECDs of the SD-type SRK receptors in insect cells was compromised by low protein yield and protein aggregation, respectively (Ma et al., 2016; Murase et al., 2020), suggesting that heterologous expression may be problematic for SD-RLKs in general. In addition, the expression of multiple protein variants in insect cells can be labour- and cost-intensive, especially if the recombinant protein yield is low.

Here, we present a versatile and simple two-step ‘depletion-binding assay’ for receptor-ligand interactions using receptor ECDs obtained in apoplastic washing fluids from *Nicotiana benthamiana* upon transient overexpression. Using the SD-RLKs LORE and SD1-23 as examples, we show that the ECDs from AtLORE and its ortholog from *Capsella rubella*, CrubLORE, bind and deplete 3-OH-C10:0 from the test solution whereas the ECD of AtSD1-23 does not. Unbound ligand was separated from receptor-ligand complexes by size exclusion filtration and detected in a sensitive bioassay using LORE-overexpressing and *lore* knockout mutant Arabidopsis plants. The depletion-binding assay developed here is of low costs and applicable for screening multiple receptor variants or ligand candidates in medium throughput. The concept can be broadly adapted for the study of receptor-ECD-ligand interactions in plants.

## RESULTS

### Receptor extracellular domains can be collected in apoplastic washing fluids upon expression in *N. benthamiana* leaves

*N. benthamiana* plants do not produce reactive oxygen species (ROS) upon 3-OH-C10:0 application, indicating they do not have a receptor for mc-3-OH-FA sensing (Fig. S1). Transient *Agrobacterium tumefaciens*-mediated expression of an AtLORE-green fluorescent protein (GFP) fusion, but not a kinase-inactive control, in *N. benthamiana* confers sensitivity to mc-3-OH-FAs (Fig. S3) (Kutschera et al., 2019). This shows that AtLORE is expressed as functional protein in *N. benthamiana*. Since AtLORE binds 3-OH-C10:0 through its ECD (Kutschera et al., 2019), the cytosolic and transmembrane domains of LORE were removed to secrete the LORE-ECD (eLORE) as soluble protein into the leaf apoplast (Fig. 1a). A red fluorescent protein, mCherry, was fused to the C-terminus of eLORE to provide a visual marker for protein localization and an epitope for immunoblot analysis (Fig. 1). As a control, mCherry was directed to the apoplast (apo-mCherry) using the signal peptide of AtLORE (amino acids 1 to 21). AtSD1-23, is the closest paralog of AtLORE, seems not to be involved in mc-3-OH-FA sensing (Fig. S3) (Ranf et al., 2015). *Capsella rubella* produces ROS upon 3-OH-FA application, suggesting is has a functional LORE receptor (Fig. S2). Upon *A. tumefaciens*-mediated expression, eAtLORE-mCherry, eCrubLORE-mCherry, eAtSD1-23-mCherry and apo-mCherry accumulated in the apoplast of *N. benthamiana* leaves and were extracted by collecting apoplastic washing fluids (AWF) (Fig. 1). The ECD-mCherry fusion proteins and the apo-mCherry control were enriched by filtration of the AWF through a membrane with 30 kDa molecular weight cut-off (MWCO) (Fig. 1).

**Figure 1.**
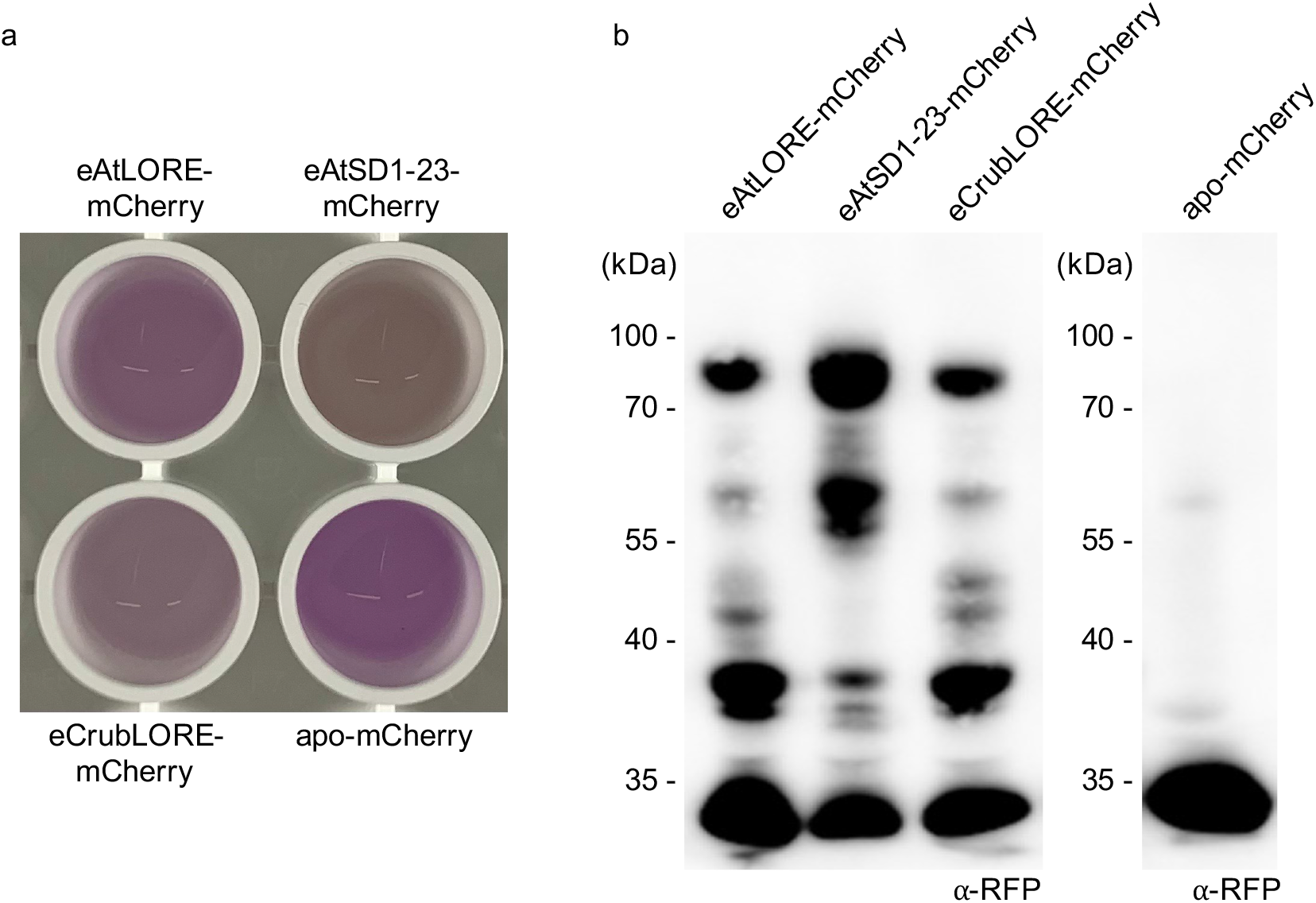
Transient expression of SD-RLK extracellular domains (ECDs) in the apoplast of *Nicotiana benthamiana* leaves. a) Photograph of apoplastic washing fluid (AWF) of *N. benthamiana* leaves expressing different SD-RLK ECD-mCherry fusion proteins and apoplastic mCherry. b) Anti-RFP immunoblot of AWF from *N. benthamiana* leaves expressing different SD-RLK ECD-mCherry fusion proteins or apoplastic mCherry (apo-mCherry). AWF corresponding to 7.5 *μ*g total proteins were loaded. Calculated molecular weights of SD-RLK ECD-mCherry fusions and mCherry are 70-75 kDa and 26.7 kDa, respectively. One representative immunoblot of three experiments is shown.

### mc-3-OH-FA depletion assay

To study the receptor-ECD-mc-3-OH-FA interaction, we exploited the molecular weight (MW) difference between proteins of interest (POI, 70-75 kDa for SD-type receptor ECD-mCherry fusions) and 3-OH-FA ligands (<1 kDa): after incubation of a defined amount of 3-OH-C10:0 ligand with concentrated AWF containing an excess of POI, the unbound ligand was separated from ECD-ligand complexes by filtration through a membrane with a MWCO of 30 kDa, which is between the MW of the free ligand and the MW of the ECD-ligand complex (Fig. 2). In case 3-OH-C10:0 binds to the POI, it is retained in the retentate (>30 kDa), whereas free 3-OH-C10:0 ligand passes through the membrane and is found in the filtrate (<30 kDa).

**Figure 2.**
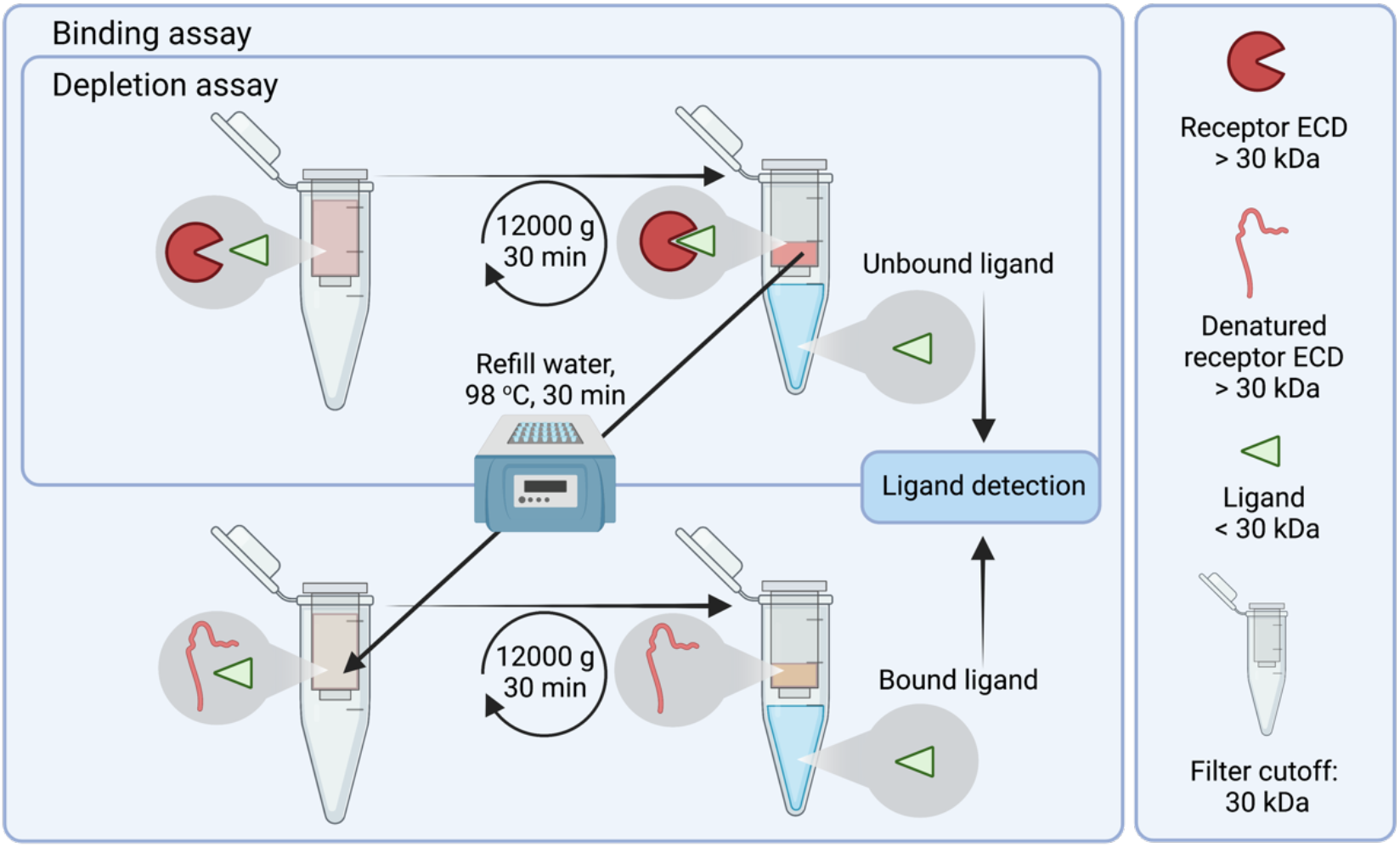
Schematic overview of depletion-binding assay. Ligand, here 3-OH-C10:0, is incubated with concentrated apoplastic washing fluids collected from *Nicotiana benthamiana* leaves transiently expressing the extracellular domain (ECD) of proteins of interest (POI) fused to mCherry. Depletion assay: The mixture is loaded into a centrifugal concentrator with a 30 kDa molecular weight cut-off. After centrifugation, unbound 3-OH-C10:0 ligand (188 Da) is found in the filtrate and POI-ECD-ligand complexes in the retentate. Unbound ligand is detected by a LORE-dependent bioassay. Depletion of the ligand from the filtrate indicates binding to the POI-ECD. Binding assay: The concentrated retentate is refilled with water up to the original volume and proteins in the retentate are heat-denatured at 98 °C for 30 minutes. The released 3-OH-C10:0 ligand is separated from the proteins by another filtration step. 3-OH-C10:0 ligand released from POI-ECD-ligand complexes is found in the filtrate and detected by a LORE-dependent bioassay. The scheme was created with BioRender.com.

Free 3-OH-C10:0 in the filtrate was detected in a sensitive bioassay using the luminescent calcium reporter aequorin to measure 3-OH-C10:0-triggered cytosolic calcium signaling in Col-0^AEQ^ Arabidopsis seedlings (Knight et al., 1991), which can robustly detect 3-OH-C10:0 at concentrations of about 50 nM and above (Fig. 3) (Kutschera et al., 2019). AWF containing apo-mCherry did not decrease 3-OH-C10:0 levels in the filtrate compared to a control containing water instead of AWF, suggesting that neither apo-mCherry nor apoplastic proteins of *N. benthamiana* bound significant amounts of 3-OH-C10:0 (Fig. 4). By contrast, 3-OH-C10:0 levels were greatly depleted in filtrates of AWF containing eAtLORE-mCherry. Thus, eAtLORE expressed in the *N. benthamiana* apoplast binds 3-OH-C10:0 (Fig. 4).

**Figure 3.**
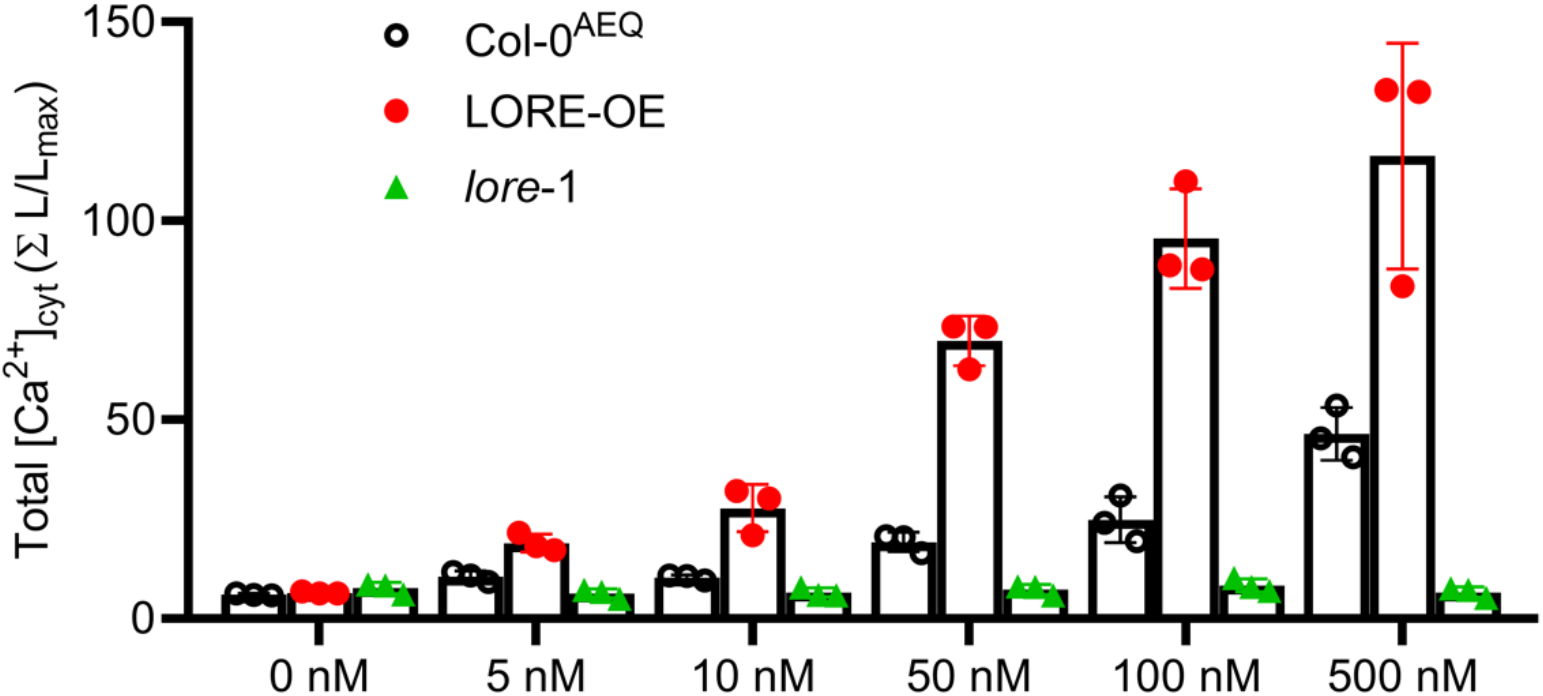
3-OH-C10:0 induces LORE-dependent increase of [Ca^2+^]_cyt_ in a dose-dependent manner. [Ca^2+^]_cyt_ measurements of Col-0^AEQ^, LORE-OE or *lore-*1 seedlings elicited with different concentrations 3-OH-C10:0 ranging from 0-500 nM (mean ± SD, n = 3 seedlings).

**Figure 4.**
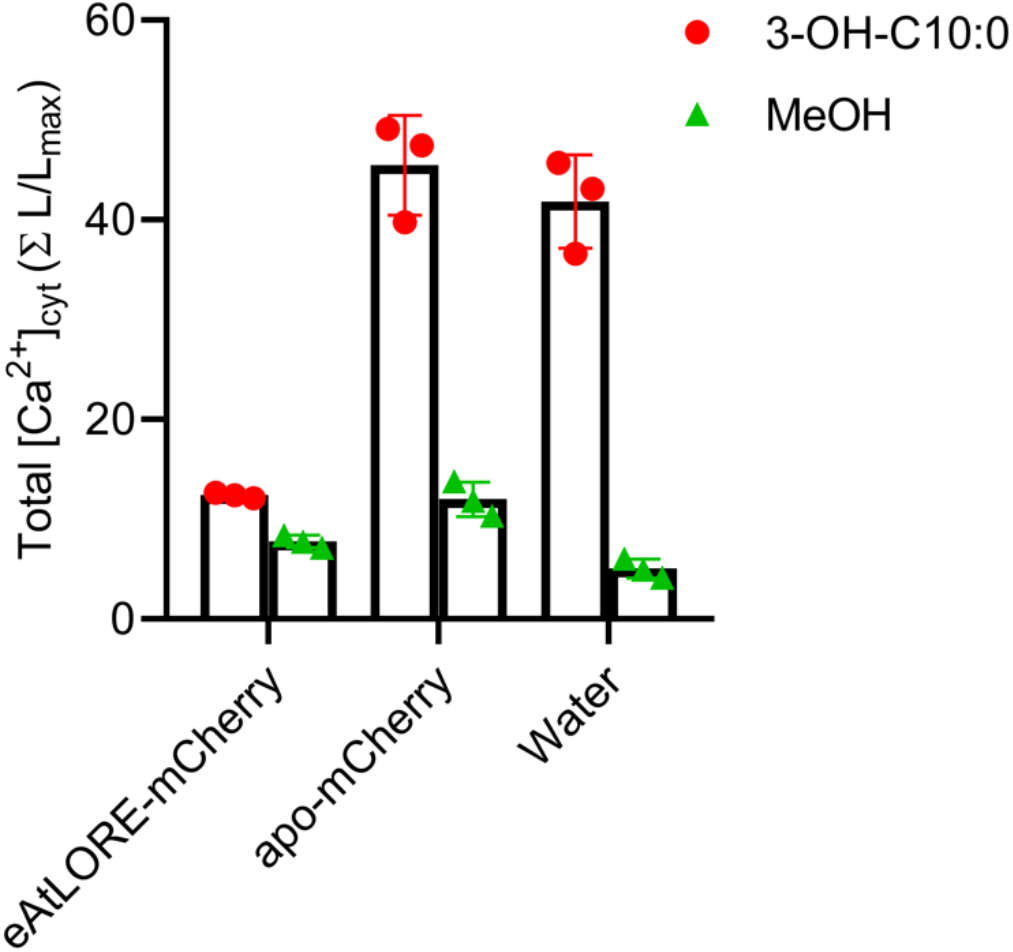
3-OH-C10:0 is depleted by eAtLORE. Total [Ca^2+^]_cyt_ levels in Col-0^AEQ^ seedlings after treatment with filtrate of depletion assay performed with apoplastic washing fluids containing eAtLORE-mCherry or apoplastic mCherry (apo-mCherry) or water control incubated with 5 µM 3-OH-C10:0 or the same volume of MeOH as control (mean ± SD, n = 3 seedlings). Total protein concentration of apoplastic washing fluids was 5.1 mg/mL. Experiment was repeated three times with similar results.

### Improved mc-3-OH-FA depletion assay using a LORE-overexpressing biosensor

Complete ligand depletion requires an excess of receptor-ECD proteins relative to the ligand concentration. To achieve this, highly concentrated AWF (total protein concentration of 5.1 mg/mL) were used in the depletion assay. Samples containing AWF+MeOH (solvent control) showed non-specific background activation in the calcium reporter assay compared to the water+MeOH control sample (Fig. 4). To reduce the amount of AWF applied in the depletion assay, we established a more sensitive plant biosensor that allows the detection of lower 3-OH-C10:0 concentrations. We generated a stable transgenic Arabidopsis line constitutively overexpressing *LORE* in the *lore*-1 mutant background, which expresses the aequorin calcium reporter (LORE-OE) (Ranf et al., 2015). LORE-OE seedlings detected 3-OH-C10:0 concentrations as low as 5 nM and responded with stronger [Ca^2+^]_cyt_ signals than Col-0^AEQ^ when treated with the same 3-OH-C10:0 concentrations (Fig. 3). With this highly sensitive biosensor, the depletion assay could be performed with a 10-fold lower 3-OH-C10:0 concentration (500 nM). While filtrates of AWF containing the apo-mCherry control activated a robust [Ca^2+^]_cyt_ signal, 70% less concentrated AWF containing eLORE-mCherry (total protein concentration of 1.5 mg/mL) was sufficient to completely deplete 3-OH-C10:0 under these assay conditions (Fig. 5). Furthermore, less concentrated AWF did not induce non-specific background activation in the calcium reporter assay in *lore*-1 mutant seedlings (Fig. 5).

**Figure 5.**
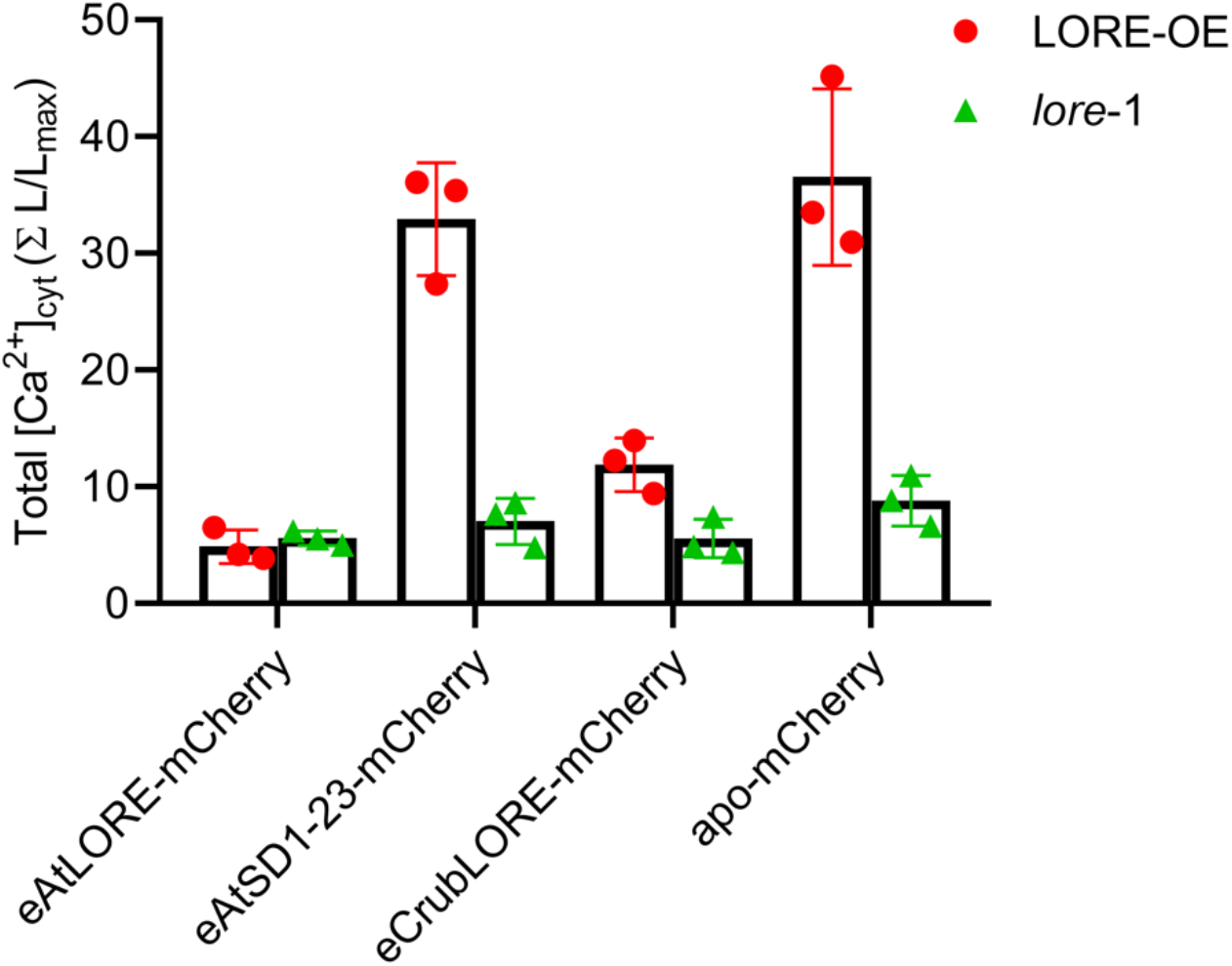
3-OH-C10:0 is depleted by AtLORE and CrubLORE extracellular domains. Depletion assay (Fig.2) was performed using 500 nM 3-OH-C10:0 and concentrated apoplastic wash fluid from *Nicotiana benthamiana* containing the indicated SD-RLK ECD-mCherry fusion proteins or apoplastic mCherry (apo-mCherry) as control. Increases in [Ca^2+^]_cyt_ were measured in Arabidopsis LORE-OE and *lore-*1 seedlings treated with the respective filtrates from the depletion assay (mean ± SD, n = 3 seedlings). Total protein concentration of apoplastic washing fluids was 1.5 mg/mL. Experiment was repeated three times with similar results.

### The LORE ortholog from *Capsella rubella* is a mc-3-OH-FA receptor

To validate the reliability of the depletion assay, we tested 3-OH-C10:0 binding by eCrubLORE-mCherry and eAtSD1-23-mCherry. 3-OH-C10:0 was depleted by AWF containing eCrubLORE-mCherry (Fig. 5), but not AWF containing eAtSD1-23-mCherry (Fig. 5). Binding of 3-OH-C10:0 to the ECD of CrubLORE demonstrates its role as mc-3-OH-FA receptor. The results also show that eAtSD1-23 cannot bind 3-OH-C10:0 and is a suitable negative control for the mc-3-OH-FA depletion assay.

### mc-3-OH-FA binding assay

The depletion assay provides indirect evidence of receptor-ECD-ligand binding by determining the proportion of 3-OH-C10:0 ligand that is not bound by a receptor protein and remains in the low MW fraction (Fig. 2). To directly measure the fraction of the 3-OH-C10:0 ligand bound by a POI, the ligand must be released from the ligand-POI complex to be available for detection by the LORE-based biosensor. 3-OH-C10:0 is stable and retains its eliciting activity even at high temperatures (Fig. S4). To release 3-OH-C10:0 from the ligand-POI complex, the proteins present in the retentate from the depletion step were denatured by heat treatment (Fig. 2). The released 3-OH-C10:0 ligand was recovered by filtration through a 30 kDa MWCO membrane and detected by the LORE-overexpressing biosensor assay (Fig. 2). The results show that eAtLORE-mCherry, but not eAtSD1-23-mCherry or apo-mCherry, binds 3-OH-C10:0 (Fig. 6). With this second binding step, the proportion of 3-OH-C10:0 ligand bound by a POI receptor can be determined directly without the need for a complex isolation procedure.

**Figure 6.**
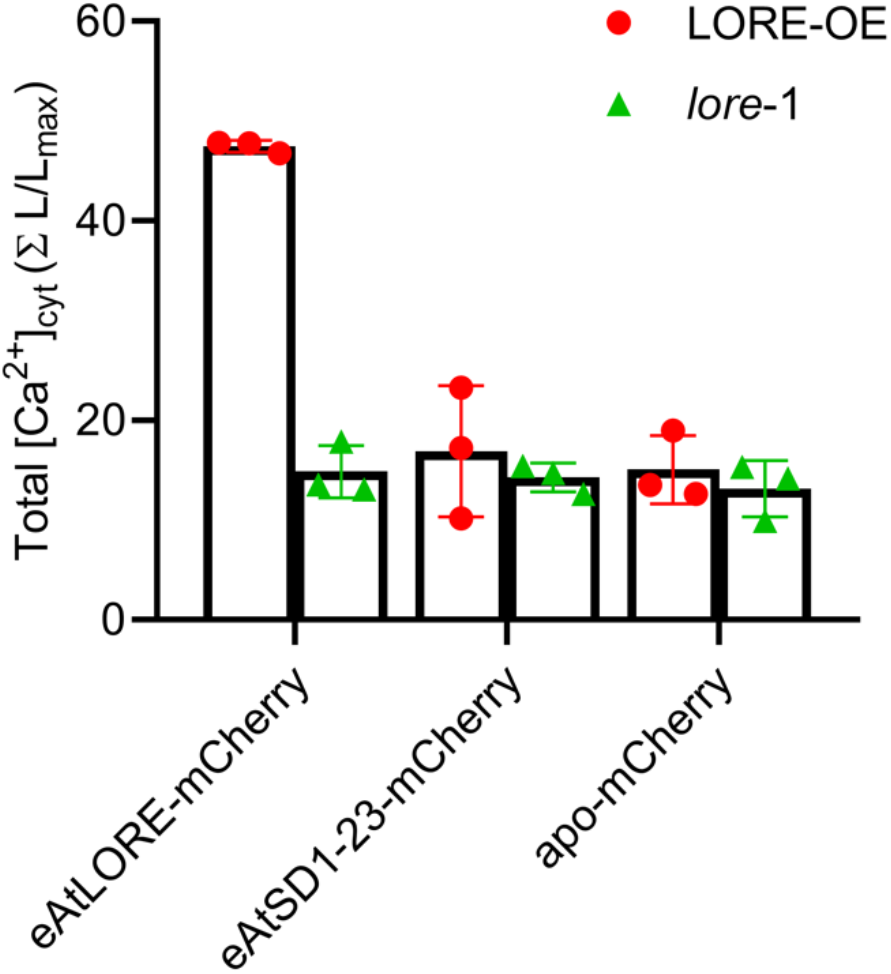
3-OH-C10:0 binds to the AtLORE extracellular domain. Binding assay (Fig. 2) was performed using 500 nM 3-OH-C10:0 and concentrated apoplastic wash fluid from *Nicotiana benthamiana* containing extracellular domains of the indicated SD-RLK ECD-mCherry fusion proteins or apoplastic mCherry (apo-mCherry) as control. Increases in [Ca^2+^]_cyt_ of Arabidopsis LORE-OE or *lore-*1 seedlings treated with the respective filtrates of heat-denatured retentates obtained in the binding assay (mean ± SD, n = 3 seedlings). Total protein concentration of apoplastic washing fluids was 1.5 mg/mL. Experiment was repeated three times with similar results.

## DISCUSSION

LORE is a PRR for mc-3-OH-FAs and its ECD binds 3-OH-C10:0 *in vitro* (Kutschera *et al*., 2019). To investigate the eLORE-mc-3-OH-FA interaction in more detail, we aimed to establish a reliable and versatile binding assay that allows screening of multiple receptor variants in medium throughput.

Several plant receptors or receptor ECDs have been successfully expressed in insect, yeast, or bacterial cells (Chinchilla et al., 2006; Liu et al., 2012; Cao et al., 2014; Choi et al., 2014). Although these widely used expression hosts can generate large amounts of recombinant protein, many problems, e.g. protein aggregation, insolubility, misfolding, misglycosylation and others, can affect the expression yield and functionality of the POI in heterologous expression systems, especially in the case of extracellular and transmembrane proteins, and make protein expression a major bottleneck for the study of receptor-ligand interactions. Heterologous expression of plant SD-type receptors seems to be a challenging task. For example, Ma et al. (2016) screened several SRK and SCR pairs for proper expression in insect cells. Expression of the ECD of wild-type SRK8 in insect cells was hampered by protein aggregation caused by patches of hydrophobic amino acids on the surface of the lectin domains. Protein engineering can help optimize heterologous expression. In case of SRK8, mutation of 11 amino acids was required to eliminate protein aggregation (Murase et al., 2020). Low yield of recombinant eAtLORE protein from insect cells and its tendency to aggregate, also made this expression system unsuitable for our purposes.

Gain of function studies in *N. benthamiana* demonstrate that it expresses functional LORE receptor protein capable of sensing mc-3-OH-FAs (Fig. S3) (Kutschera et al., 2019). Thus, *N. benthamiana*, lacking a constitutive 3-OH-FA receptor (Fig. S1), is a suitable host to produce functional SD receptor protein that can be used for 3-OH-FA binding assay without an elaborate purification procedure. Since the ECD is the mc-3-OH-FA-binding domain of LORE (Kutschera et al., 2019), we expressed only the POI-ECDs, which can be easily collected from *N. benthamiana* leaves in AWF. The receptor ECDs have to pass the intrinsic protein quality control system of the host secretory pathway and can be assumed to be correctly folded and post-translationally modified. This plant expression method yields receptor ECDs in soluble form without the need to generate cell lysates and solubilize membranes, which could interfere with subsequent binding studies. Although the yield of POI-ECD in *N. benthamiana* was only moderate and we observed partial degradation in immunoblot analysis (Fig. 1b), the amount of POI-ECD recovered in AWF was sufficient for the depletion-binding assay after concentration, presumably because it was used in excess relative to the ligand. Whereas strongly concentrated AWF caused some unspecific background responses during the detection of the 3-OH-C10:0 ligand in the calcium signaling bioassay using Col-0^AEQ^ seedlings (Fig. 4), this was not observed when performing the depletion assay with less concentrated AWF and using the more sensitive LORE-OE biosensor (Fig. 5).

Since covalent modifications of the 3-OH-C10:0 ligand affect its eliciting activity (Kutschera et al., 2019), presumably by impairing receptor binding, fluorescence labelling of mc-3-OH-FAs or cross-linking to a resin are not possible. The depletion-binding assay was therefore designed to study interaction of the POI with unlabelled 3-OH-C10:0 ligand in solution (Fig. 2). We used a highly sensitive LORE-OE plant line as biosensor to detect 3-OH-C10:0 at nanomolar concentrations (Fig. 3). However, the use of a LORE-dependent biosensor restricts detection to potential ligands that activate LORE signalling, at least in the depletion assay. For example, the LORE-OE biosensor cannot be used to investigate depletion of 3-OH-FAs with longer acyl chains, such as 3-OH-C14:0, or modifications of the 3-OH-group (Kutschera et al., 2019). Binding of such ligands to eLORE could be tested indirectly in the binding step by outcompeting binding of 3-OH-C10:0. Alternatively, ligands could be detected by other analytical means, such as high-performance liquid chromatography-coupled mass spectrometry, which is however more labour-intensive and requires specialized instrumentation or facilities.

The results of the depletion-binding assay obtained here with eAtLORE are consistent with previous findings (Kutschera et al., 2019) and further demonstrate a direct eAtLORE-3-OH-C10:0 interaction. To further test the reliability and applicability of the depletion-binding assay, we tested the ECDs of two additional SD-RLKs, eAtSD1-23 and eCrubLORE, for binding of 3-OH-C10:0, that share 81% and 92% amino acid identity with eAtLORE, respectively. eAtSD1-23 neither depleted nor bound detectable amounts of 3-OH-C10:0. *C. rubella*, a close relative of *A. thaliana*, is sensitive to mc-3-OH-FAs (Fig. S2). Its putative LORE ortholog, CrubLORE, was identified here as a new mc-3-OH-FA-binding receptor. These results will help to better understand mc-3-OH-FA sensing by LORE in the future.

From protein expression to interaction assay, the depletion-binding assay developed here for investigating interactions between plant-expressed SD-type receptor ectodomains and mc-3-OH-FAs relies on simple techniques and routine lab equipment available in most plant research labs. The workflow of the assay is very flexible with regard to the protein source and ligand detection method and can be easily adapted to study other receptor-ligand pairs, especially low-molecular weight ligands. The easy workflow makes this assay applicable for screening receptor-ligand interactions in medium throughput.

## MATERIALS AND METHODS

### Plant material

*A. thaliana* Col-0 expressing pCaMV35S:apoaequorin in the cytosol (Col-0^AEQ^) was obtained from M. Knight and the *lore*-1 mutant was described previously (Ranf et al., 2015). *C. rubella* (N22697) was obtained from the European Arabidopsis Stock Centre. After stratification at 4°C in the dark for three days, *C. rubella* was grown on soil with vermiculite (9:1) in a climate chamber with 8 hours of daylight at 22°C/20°C (day/night) in 55–60% relative humidity. *N. benthamiana* was grown on soil with vermiculite (9:1) in a climate chamber with 16 hours of daylight at 23°C/21°C (day/night) in 55–60% relative humidity.

### ROS detection in *N. benthamiana* and *C. rubella* leaves

ROS production in leaves of 6 weeks-old *N. benthamiana* and *C. rubella* after elicitation by 3-OH-C10:0 (Matreya LLC) and flg22 (QRLSTGSRINSAKDDAAGLQIA) was measured as described (Ranf et al., 2015). Leaf discs were incubated on water overnight. Shortly before the measurements, water was exchanged for 100 µL water with 5 µM L-012 (WAKO chemicals) and 2 µg/mL horseradish peroxidase (Type II, Roche). After the leaf discs were treated with 1 µM 3-OH-C10:0 (stock in MeOH) or 500 nM flg22 (stock in water), accumulation of ROS was monitored for 45 min in a microplate luminometer (Infinite F200 PRO, Tecan, or Luminoskan Ascent 2.1, Thermo Scientific). MeOH and water were used as control treatments. Total luminescence signals were calculated by summing up relative light unit (RLU) from 1-45 min.

### Molecular cloning

Cloning of At*LORE* (*SD1-29*, At1g61380) was described previously (Ranf et al., 2015). Total RNA from *A. thaliana* and *C. rubella* leaf material was extracted by the Trizol method, treated with DNase I (Thermo Scientific) and reverse-transcribed using oligo(dT)_18_ and RevertAid reverse transcriptase (Thermo Scientific) according to the manufacturers’ instructions. *A. thaliana* At*SD1-23* (At1g61390) and *C. rubella* Crub*LORE* (CARUB_v10021901mg) were amplified from cDNA with primers having BpiI recognition sites and cloned into a modified GoldenGate-compatible pUC18 vector (Ranf et al., 2015). Internal BpiI, Esp3I and BsaI recognition sites were curated by site-directed mutagenesis. An additional intron (1^st^ intron from *LORE*, 109 bp) was inserted into At*SD1-23* at the 65^th^ codon. Sequences of At*LORE*, At*SD1-23* and Crub*LORE* corresponding to the ECDs were amplified from cloned genes with primers having BpiI sites and inserted into same modified GoldenGate-compatible pUC18 vector. The coding sequences were assembled with a CaMV35S promoter and terminator with or without a C-terminal ten-glycine linker and an mCherry or GFP epitope tag. The sequence corresponding to the signal peptide (amino acids 1-21) of LORE was fused to the N-terminus of mCherry for secretion into the apoplast. Primers used for cloning are listed in Supplementary table 1. Sequences of all cloned expression cassettes were verified by Sanger Sequencing. Expression cassettes were moved into a modified GoldenGate-compatible *Agrobacterium tumefaciens* binary vector pCB302 by GoldenGate techniques and transformed into *A. tumefaciens* GV3101 for generation of transgenic Arabidopsis lines and protein expression in *N. benthamiana*.

### Generation of LORE-overexpression line

LORE overexpression line (LORE-OE) was generated using the floral dip method (Logemann et al., 2006). *lore*-1 was transformed by *A. tumefaciens* GV3101 carrying pCB302-pCaMV35S:AtLORE. Dipped plants were grown under long day conditions and mature seeds were harvested. Transformed plants having BASTA resistance were identified by glufosinate ammonium selection and phenotypically selected for LORE overexpression by screening for a strong 3-OH-C10:0-activated calcium signaling response. Homozygous plants (T_3_ and above) were used in all assays.

### Cytosolic Ca^2+^ measurement

Seed sterilization, growth of seedling, and calcium measurement were described previously (Ranf et al., 2012). Apoaequorin-expressing seedlings cultured in liquid 0.5 × Murashige & Skoog medium for 9 days were transferred into 96-well plates and incubated in 100 µL water with 10 µM coelenterazine-h (p.j.k. GmbH) per well overnight. Luminescence signal was measured in a microplate luminometer (Infinite F200 PRO, Tecan, or Luminoskan Ascent 2.1, Thermo Scientific) with a 10 sec interval. Each seedling was treated with 25 µL of test sample. Luminescence signals (relative light units, RLU) were converted to L/Lmax (luminescence counts per sec/total luminescence counts) as described (Ranf et al., 2012). Total calcium signals were calculated by summing up L/Lmax from 2-30 min.

### Transient protein expression in *N. benthamiana* leaves

Proteins of interest were expressed in leaves of 6-7 weeks-old *N. benthamiana* plants via Agrobacterium-mediated transformation. *A. tumefaciens* GV3101 carrying different expression constructs were grown on LB agar with 30 µg/mL gentamycin, 10 µg/mL rifampicin, and 50 µg/mL kanamycin for 48 hours, and then inoculated and cultured in AB medium supplemented with 100 µM acetosyringone and 50 µg/mL kanamycin at 28°C, 200 rpm overnight. Agrobacteria were harvested (1900 g, 10 min), washed three times with infiltration buffer (10 mM MgSO4, 10 mM MES pH 5.5, 200 µM acetosyringone), and incubated at 28°C for three hours. The OD_600_ of Agrobacteria was measured and adjusted to 0.5 with infiltration solution. All Agrobacteria clones were mixed with an equal volume of Agrobacterium carrying the silencing suppressor p19. Agrobacteria mixtures were infiltrated into fully expanded leaves of *N. benthamiana* using a needleless syringe.

For analysis of ROS production, Agrobacteria carrying *LORE-GFP, LORE-Km-GFP*, and *SD1-23-GFP*, were cultured and harvested as described above. The OD_600_ of Agrobacterium was adjusted to 0.025, and Agrobacterium carrying constructs were mixed 1:1 with Agrobacterium carrying the p19 silencing suppressor. For ROS detection, leaf discs from infiltrated leaves were harvested 36 hours after infiltration and recovered for 6 hours on water before the experiment.

### Immunoblot analysis

5 *μ*L of concentrated AWFs were separated by 10% SDS-PAGE, and transferred onto a nitrocellulose membrane using a Semi-Dry Transfer Cell (Bio-Rad). The membrane was blocked using BlueBlock PF (SERVA) and developed using rat monoclonal anti-RFP 5F8 (1:2000, Chromotek) and rabbit anti-rat-peroxidase A9542 (1:20000, Sigma-Aldrich) with SuperSignal West Dura Extended Duration Substrate (Thermo Scientific).

### 3-OH-C10:0 depletion-binding assay

Apoplastic wash fluids (AWFs) were harvested from leaves of *N. benthamiana* 5 days after agrobacterium-mediated transformation as described before with minor modifications (Rutter et al., 2017). Leaves were vacuum-infiltrated with water and AWF were collected by centrifugation for 20 minutes, at 800 g and 4°C. AWF were cleared by centrifugation (20 min, 8000 *g*, 4^°^C), filter-sterilized (pore size 0.22 μm), desalted (PD-10 Desalting Columns, Cytiva), and concentrated to the indicated total protein concentration (Vivaspin 20, 30 000 MW cut-off, Sartorius). For depletion assay, AWF concentrate was mixed in 9:1 ratio (v:v) with the indicated concentration of synthetic 3-OH-C10:0 and incubated for one hour at 4^°^C on a rotator. Unbound ligands were separated from the mixture (Vivaspin 500, 30 000 MW cut-off, Sartorius). For binding assay, the retentate was refilled with water to the original volume of the AWF-3-OH-C10:0 mixture described above, transferred to a new tube and incubated at 98°C for 30 minutes. The bound ligand released from heat-denatured AWF was isolated by Vivaspin 500 concentrators with 30000 MW cutoff. Content of unbound 3-OH-C10:0 in filtrates was assessed by [Ca^2+^]_cyt_ measurements using Col-0^AEQ^, LORE-OE and *lore*-1 seedlings as indicated with a filtrate:water ratio of 1:4 (v:v).

## ACKNOWLEDGEMENTS

Work in the Ranf lab is supported by the German Research Foundation (SFB924/TP-B10 and Emmy Noether programme RA2541/1).

## SUPPLEMENTARY FIGURES AND TABLES

Supplementary figures S1-S4 and supplementary table 1 are provided in separate files.

