## Supplementale Table S1 for "Low cost, medium throughput depletion-binding assay for screening S-domain-receptor ligand interactions using *in planta* protein expression"

### Supplementary Table 1. Primers used in this study

| Target (Gene ID) | Primer | Sequence (5’ → 3’) | Purpose |
|---|---|---|---|
| AtLORE (At1G61380) | SD129-START | tcgtctctAATGGGTATGGTTTTATTTGCTTGC | Cloning LORE ECD |
| AtLORE (At1G61380) | SD129-ED-R | tgaagacttaCTAGCTTCCAGCCAATTCTG | Cloning LORE ECD and CrubLORE ECD |
| CrubLORE (CARUB_v100219 01mg) | CARUB29- START | tgaagacttAATGGGTATGGTTTTATTTGC | Cloning CrubLORE |
| CrubLORE (CARUB_v100219 01mg) | CARUB29-EspF | tgaagactcCGCCTGGGGAGTTCACAC | Cloning CrubLORE |
| CrubLORE (CARUB_v100219 01mg) | CARUB29-EspR | tgaagacttGGCGACGGATCACTGTAAC | Cloning CrubLORE |
| CrubLORE (CARUB_v100219 01mg) | CARUB-STOPm | tgaagacttACTAGCGTCCTTGGATCATAGATTG | Cloning CrubLORE |
| mCherry | apomCh_F | tttcgtctctAATGGTGAGCAAGGGCGAGG | Cloning apoplastic mCherry with AtLORE signal peptide |
| mCherry | apomCh_R | tttcgtctctCATTGCATAGCCACAAGTTGG | Cloning apoplastic mCherry with AtLORE signal peptide |
| AtSD1-23 (At1g61390) | SD123_START | tgaagacttAATGTACAAACTTCCACAAAG | Cloning AtSD1-23 |
| AtSD1-23 (At1g61390) | SD123-BpiM_R | tgaagacgagGACTGAGGCAGCATAGTATTACC | Cloning AtSD1-23 |
| AtSD1-23 (At1g61390) | SD123-BpiM_F | tgaagaccaGTCCTCTGTGATGTATGATATTCC | Cloning AtSD1-23 |
| AtSD1-23 (At1g61390) | SD123-EspM_R | tgaagactcTGACACACCCACTTGTCCAATTC | Cloning AtSD1-23 |
| AtSD1-23 (At1g61390) | SD123-EspM_F | tgaagacgtGTCAGACGTACACAATTATC | Cloning AtSD1-23 |
| AtSD1-23 (At1g61390) | SD123-BsaM_R | tgaagaccaGATCTCGGTGAATTACCC | Cloning AtSD1-23 |
| AtSD1-23 (At1g61390) | SD123-BsaM_F | tgaagacgaGATCTGAAGGTCAGCAAC | Cloning AtSD1-23 |
| AtSD1-23 (At1g61390) | SD123-STOP | tgaagacttACTAACGCCCTTGAATCAC | Cloning AtSD1-23 |
| AtSD1-23 (At1g61390) | SD123intron_F | tttgaagactcAGCTTCTTCAGTCCTAATAATTC | Cloning AtSD1-23, intron insertion |
| AtSD1-23 (At1g61390) | SD123intron_R | tttgaagacttACCCTAGCTCATA AACTCC | Cloning AtSD1-23, intron insertion |
| AtLORE (At1G61380) | 129intron-F | tttgaagacttGGGTAAGGATAAAATACATTTCTTCC | Cloning AtSD1-23, intron insertion |
| AtLORE (At1G61380) | 129intron-R | tttgaagacgaAGCTGAGATATTCACCAG | Cloning AtSD1-23, intron insertion |
| AtSD1-23 (At1g61390) | SD123-EC-R | tgaagacttaCTAGCTACCAGCCAATTCTG | Cloning AtSD1-23 ECD |
