## Supplemental Figures S1-S4 for "Low cost, medium throughput depletion-binding assay for screening S-domain-receptor ligand interactions using *in planta* protein expression"

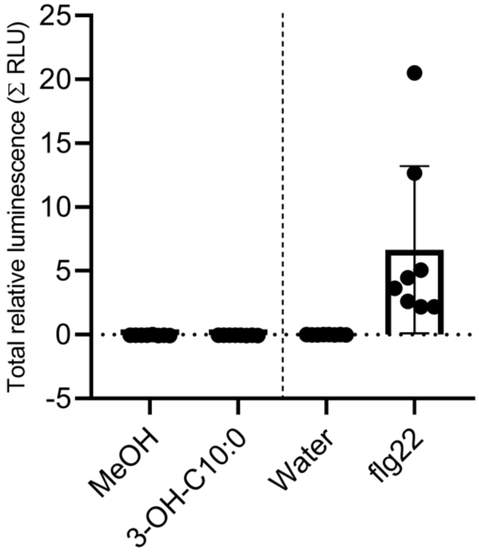


**Figure S1.** ***Nicotiana benthamiana*** **does not produce ROS in response to 3-OH-C10:0**. Leaf discs of *N. benthamiana* were treated with 1 $\mu$M 3-OH-C10:0, MeOH (solvent control of 3-OH-C10:0), 0.5 $\mu$M flg22 or water (control) (mean ± SE, n = 8). Luminescence (relative light units, RLU) was measured at one-minute intervals over a 45-minute period. RLU of treated samples were summed up after subtraction of RLU of water control.

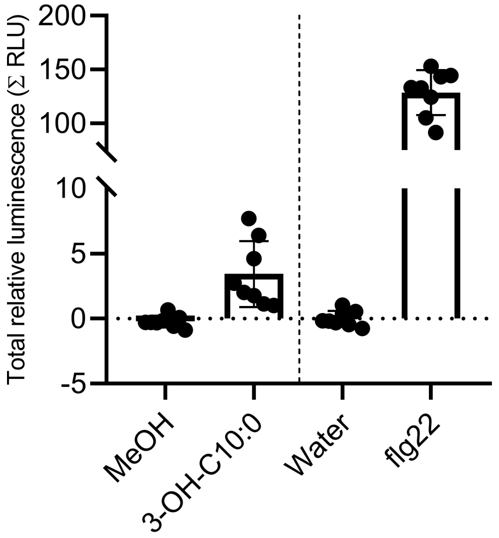


**Figure S2. *Capsella rubella* produces ROS in response to 3-OH-C10:0*.*** Leaf discs of *C. rubella* were treated with 1 $\mu$M 3-OH-C10:0, MeOH (control of 3-OH-C10:0), 0.5 $\mu$M flg22 or water (control) (mean ± SE, n = 8). Luminescence (relative light units, RLU) was measured at one-minute intervals over a 45-minute period. RLU of treated samples were summed up after subtraction of RLU of water control.

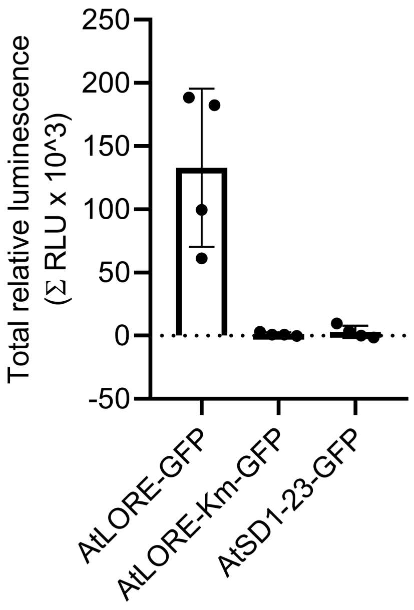


**Figure S3. Transient expression of AtSD1-23 in *Nicotiana benthamiana* does not confer sensitivity to 3-OH-C10:0**. ROS accumulation in leaf discs of *N. benthamiana* expressing AtLORE-GFP, kinase-inactive AtLORE-Km-GFP and AtSD1-23-GFP elicited with 1 $\mu$M 3-OH-C10:0 or the same amount of MeOH (control) was measured (mean ± SE, n = 4). Luminescence (relative light units, RLU) was measured at one-minute intervals over a 45-minute period. RLU of treated samples were summed up after subtraction of RLU of MeOH control.

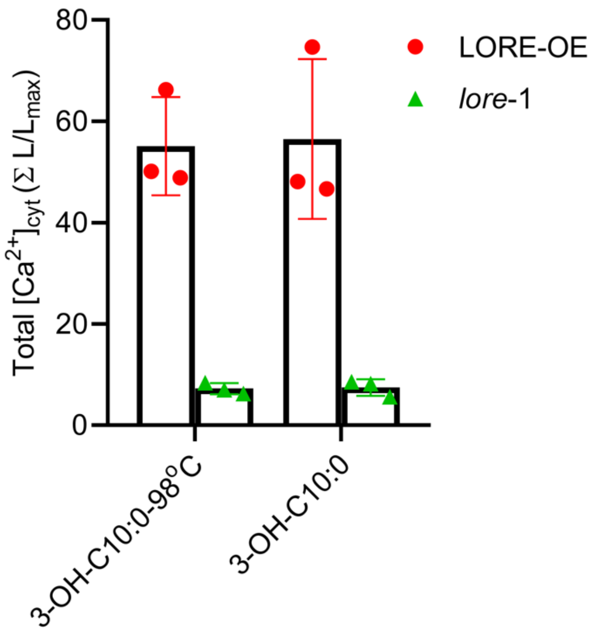


**Figure S4. Elicitor activity of 3-OH-C10:0 after heat treatment.** $[Ca^{2+}]_{cyt}$ was measured in *Arabidopsis* seedlings of LORE-overexpressing line (LORE-OE, mean ± SD, n = 3 seedlings) and *lore*-1 (mean ± SD, n = 3 seedlings) treated with 100 nM 3-OH-C10:0 or 500 nM 3-OH-C10:0 heat-treated at 98 $^{o}$C for 30 minutes.
